# ChemoTrack: A comprehensive dataset linking single-cell migration trajectories to precisely defined chemotactic signals

**DOI:** 10.64898/2026.06.28.734951

**Authors:** Devi Prasad Panigrahi, Naoto Sakurai, Lucija Mijanovic, David M. Versluis, Luke Tweedy, Philip Pearce, Laura M. Machesky, Robert H Insall

## Abstract

Chemotaxis drives cell migration in processes ranging from wound healing to embryonic development and cancer metastasis, yet its quantitative understanding remains limited because responding cells change and degrade attractant gradients, and existing datasets are too small and imprecise to capture stochastic behaviour. We present ChemoTrack, a publicly accessible resource comprising 2 million measurements from 500,000 migration tracks, in which the chemoattractant gradient and concentration experienced by every cell at the time of observation are precisely determined. The dataset includes microscopy images and trajectories spanning a full range of biologically relevant chemotactic conditions. Analysis shows that cells steer according to absolute differences in active receptor number, not fractional receptor occupancy, and maximal chemotaxis is not predicted by half-maximal receptor occupancy. By combining scale, precision and accessibility, ChemoTrack shifts quantitative description of eukaryotic chemotaxis from experimental conditions to the instantaneous chemical signal experienced by individual cells, enabling future mathematical and mechanistic analyses.

## I. INTRODUCTION

Chemotaxis, the directed migration of cells in response to chemical signals, is a widespread and evolutionarily conserved mechanism that steers cell motility and thus underpins diverse areas of physiology. The attractant chemicals - chemoattractants - are dissolved in the external fluid environment of the cells, and are sensed by transmembrane receptors that become activated upon binding to a chemoattractant molecule. Although the molecular mechanisms governing bacterial chemotaxis are well characterized [1], a similar level of understanding has yet to emerge for the chemotaxis of eukaryotic cells. It is clear that eukaryotic chemotaxis plays a key role in biological processes ranging from mammalian embryonic development [2] and immune responses [3], to unicellular feeding [4], and pathological processes like pathogen invasion [5] and cancer metastasis [6]. Across all these physiological contexts, chemotaxis drives directional steering during cell migration. For example, during embryonic development the spatial organization of neurons in the brain is governed by cell migration through a pattern of chemoattractants [7], and impaired chemoattractant-receptor activity in embryos has been found to result in defective brain development [8].

True understanding of the mechanism of chemotaxis will clearly require a fully quantitative understanding of the details of steering. Although there has been some work on understanding the roles of different proteins in the chemotactic pathway [9, 10], a precise quantification of chemotactic sensing is lacking in the literature. While in vitro experiments allow control over the chemoattractant gradients that are initially applied, cells are not passive responders, and have long been known to greatly modify what they encounter. They may modify or even create [11, 12] gradients by breaking down the attractants they encounter or by removing them by receptor-mediated endocytosis [13]. More strikingly, they may complicate their environments by secreting their own secondary chemoattractants [14], or chemorepellents [5], often under the direct control of the applied attractants [15], thereby creating spatially and temporally distinct gradients that convolve measurements. This may even lead to robust self-generated cues in the absence of any imposed gradient [16–19]. Even though the effects of chemoattractant secretion can be restricted using microfluidic devices [20–22], the effects of chemoattractant degradation cannot be avoided because it occurs at length scales that are comparable to receptor-ligand binding [23].

As a result of this, most existing quantitative data on chemotaxis of cells towards a native chemoattractant, for example, chemotaxis of *Dictyostelium discoideum* cells towards cAMP [20, 21, 24], typically capture the combined effects of chemotactic sensing, chemoattractant degradation, and secondary attractant production. They cannot therefore be considered as data quantifying chemotactic sensing alone. Degradation and new attractant production are extremely difficult to measure locally in real time. Although these limitations can be partially addressed using photo-activated chemoattractants and microfluidic devices, the amount of data generated from these assays is limited [25]. Since chemotactic response of cells is highly stochastic, and chemotactic migration has been found to emerge from an integration of multiple decision-making processes accumulated over time [22, 26, 27], understanding the mechanisms underlying chemotactic migration requires a statistical analysis of a large number of quantitatively consistent measurements that are made on the timescale of decision-making by an individual cell. To our knowledge, there is currently no dataset containing individual cell tracks measured under precisely calibrated chemoattractant concentrations and gradients. The absence of quantitative data on chemotactic sensing not only limits mechanistic understanding of chemotaxis but also hinders the development of detailed mathematical models for cell migration in complex environments, such as a combination of chemical signals and complex geometry [28].

In this work we have created a general dataset for chemotactic sensing in eukaryotic cells. Mammalian cells have complex signalling pathways which have not been described fully, with high levels of autocrine chemoattractant signalling that cannot currently be accurately measured, making it impossible with current technology and understanding to describe the chemotactic environment of a cell precisely. To address this, we used *Dictyostelium discoideum* cells that chemotax using an evolutionarily-conserved G-protein-coupled receptor mechanism, which uses the same mechanism in detail as mammalian cells responding to chemoattractants such as FMLP and IL-8 [29]. *D. discoideum* cells detect cycylic AMP (cAMP) through the G-protein-coupled receptor cAR1, a member of the calcitonin receptor family whose pharmacology is understood in great detail. This has made *D. discoideum* a powerful system for investigating fundamental principles of eukaryotic chemotactic sensing [30]. However, the cells secrete their own, autocrine cAMP, which substantially alters the chemoattractant field. In our experience, cells secrete significant cAMP even in the presence of inhibitors like caffeine, complicating measurement of cells’ local environments. To mitigate this, we have used cells that have been genetically modified by knocking out [31] the acaA gene that encodes the endogenous secretion of cAMP in response to starvation [32], giving cells that do not secrete their own cAMP.

In addition, the chemoattractant cAMP is subject to degradation by the cells. To circumvent this, we employed the chemically modified analog Sp-cAMPS, which binds to and activates the cAR1 receptor but is resistant to degradation by cell-secreted phosphodiesterases. This resistance results from a structural modification in Sp-cAMPS in which one non-bridging oxygen atom of the phosphate group of cAMP is substituted with sulfur. Moreover, Sp-cAMPS has a relatively low binding affinity to cAR1 as compared to cAMP, thereby reducing the effects of alteration in chemotactic sensing due to receptor endocytosis [33]. To obtain a large dataset of cells exposed to identical chemotaxis conditions, we employed direct-view chemotaxis chambers for visualizing cell migration [34] along a domain having a one-dimensional gradient of Sp-cAMPS. Taken together, this gives us a system where we have a large ensemble of cells that are exposed to a precisely defined gradient and absolute chemoattractant concentration everywhere in the domain of interest.

We present ChemoTrack, a dataset comprising 32 different chemotactic gradients, along with 8 control conditions with no gradients but different backgrounds, each consisting of approximately 14000 cell tracks on an average. We identified an appropriate metric for quantifying chemotactic sensing based on the existing literature [35], and a comparison of how other commonly used metrics perform for our dataset. Based on this, we computed chemotactic sensing in a statistically consistent manner and found that the conditions provided in ChemoTrack are sufficient to capture the biologically relevant regime for chemotactic sensing. We found that in contrast to the classical literature on chemotactic sensing [36], the chemoattractant concentration for optimal chemotactic response is not necessarily equal to the dissociation constant for the receptor-ligand binding. Furthermore, we assessed the differences in chemotactic sensing arising from cellular chemoattractant degradation and self-secretion, demonstrating that these processes can substantially alter the apparent chemotactic sensitivity measured in experiments. These findings show that quantitative measurements of chemotaxis reported in the literature can be significantly influenced by cell-mediated modification of the imposed chemoattractant field, highlighting the importance of our experimental pipeline and dataset for quantitative characterization of chemotactic sensing.

The ChemoTrack dataset generated during this study has been deposited in the BioImage Archive under accession number (S-BIAD3674), and is publicly available. The size of the dataset, along with precise knowledge of the chemoattractant concentrations and gradients corresponding to every data point, provides a detailed summary of quantitative knowledge about chemotactic sensitivity which will be crucial for gaining insights into mechanisms for chemotactic sensing.

## RESULTS

### Creating an extensive and fully quantitative chemotaxis dataset

We examined a strain derived from wild type *Dictyostelium discoideum* NC4 cells, grown using bacteria as a food source, to avoid the complicating effects on cell migration caused by axenic growth and loss of the NF1 RasGAP [37]. Additionally, we find that this gives cells whose morphology and behavior are consistently more homogeneous and consistent than cells grown axenically [38]. Use of an acaA^−^ mutant [31] ensured that cells were unable to distort results by secreting their own cAMP (see validation below). Because wild-type cells use autocrine cAMP signaling to drive differentiation, we differentiated the cells using exogenous cAMP pulses (see Methods for details). This approach has the added advantage of minimizing heterogeneity in chemotactic ability of cells within the population, and making cells differentiated on different days behave consistently with one another. The cells were then introduced into direct-view Insall chemotaxis chambers [34], and exposed to gradients of the chemoattractant Sp-cAMPS that cannot be degraded by the cells [39] (see Methods for more details). Additionally, Sp-cAMPS has a far lower affinity than cAMP for cAR1, so we use approximately 1000x more, greatly diminishing the cells’ ability to deplete the Sp-cAMPS by endocytosis. Together, these give us a system where we can expose a population of cells to a tightly-defined, one-dimensional linear chemoattractant gradient, and the chemoattractant concentration can be calculated everywhere in the domain with the knowledge of the chemoattractant concentrations at the boundaries of the viewing bridge (see Fig. 1A).

**FIG. 1.**
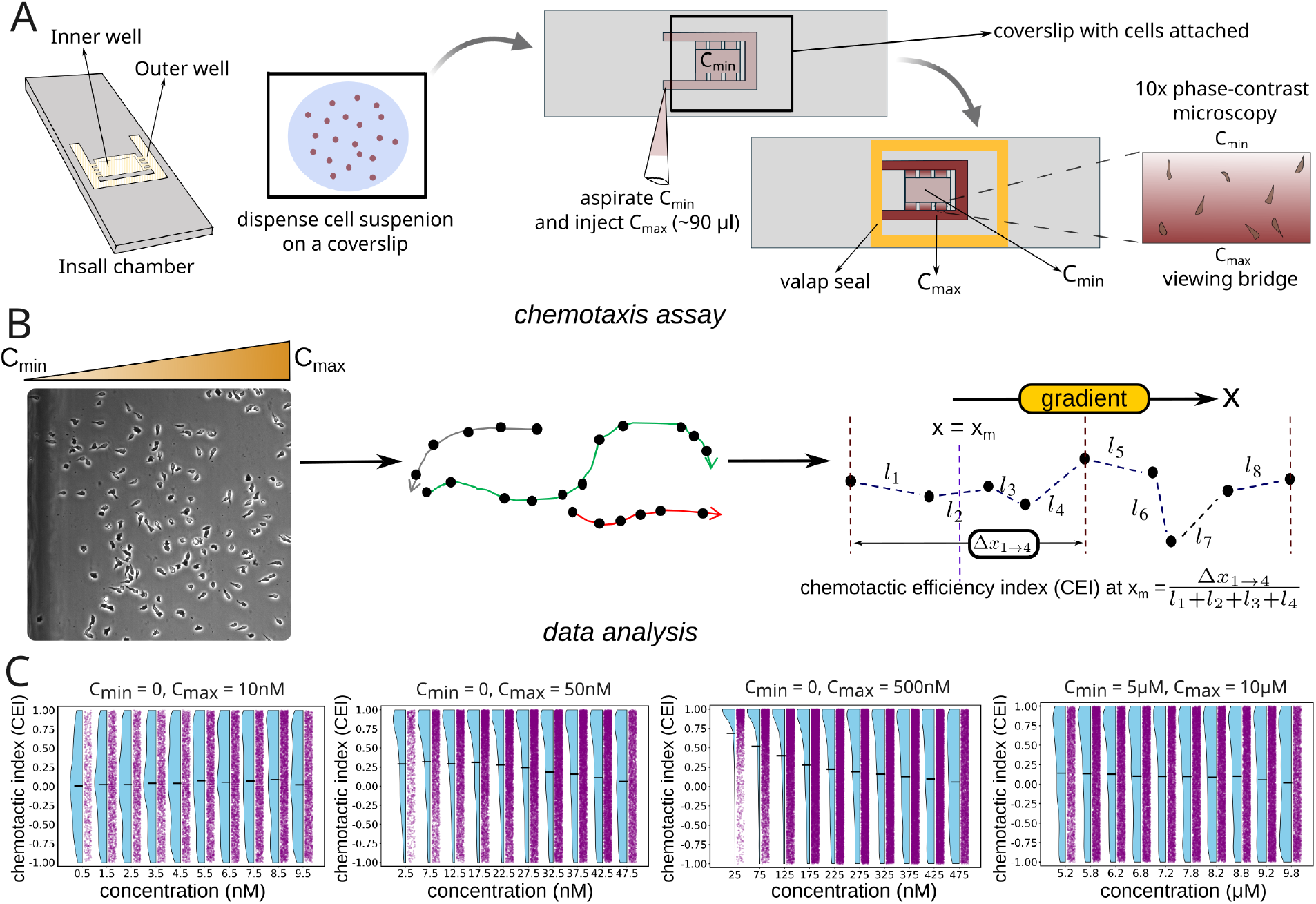
ChemoTrack: pipeline and processing of cell tracks. (A) Schematic of the Insall chamber showing the geometry and assembly of the chemotaxis chambers. A chemoattractant gradient was established between the inner and outer wells and cells were imaged on the viewing bridges. (B) Schematic of the cell tracking and analysis: cells were segmented using Fiji and tracked using an in-house Fiji plugin. Chemotactic sensing was quantified using the chemotactic efficiency index, only for those segments which are longer than 15 *µm*. (C) Representative examples for dose-response of chemotactic sensing with respect to chemoattractant concentration. Each panel shows a gradient with the start and end concentrations in the label, aggregated from 3 independent experiments. Every CEI measurement is associated with a chemoattractant concentration, and all measurements from a chemotaxis condition are are divided into 10 bins uniformly distributed between C_min_ and C_max_, and the mean chemoattractant concentration corresponding to each bin is shown on the x-axis. For each concentration bin, individual CEI measurements are shown as purple dots on the right, the overall distribution is represented by a blue half-violin plot on the left, and the mean CEI is indicated by a horizontal black line.

Cells were followed using time-lapse microscopy, and imaged at 30 second intervals. The resulting images were processed using Fiji [40], with cells segmented and tracked using an in-house Fiji plugin (see Methods for details). Cell tracks can potentially span the entire width of the domain along which the gradient is imposed, and therefore, cells experience a different chemoattractant concentration at every point on the track. Since the imposed chemoattractant gradient is constant in both space and time, and the concentration is known at every point in the domain, we can compute the chemotactic efficiency of cells across all background concentrations. To do this, we spliced the track into segments that capture the motion of a cell in an interval of two minutes, containing five data points for the cell position. We used the chemotactic efficiency index (CEI) as the metric for quantifying chemotaxis [35] as we found it to be the more robust as compared to other commonly used chemotaxis metrics (see Supplementary Materials). CEI is defined as the ratio of the distance moved by a cell along the direction of the imposed gradient, divided by the total distance traversed by the cell during the course of observation (see Fig. 1B). The CEI corresponding to the midpoint of the segment is defined as the ratio between the displacement of the cell along the direction of the imposed gradient (Δ*x*_1→4_), and the total distance covered by the cell in that segment (*l*_1_ + *l*_2_ + *l*_3_ + *l*_4_).

This ensures that the cell displacement over the timescale of one measurement is not large enough to incur significant error in the background chemoattractant concentration, and the chemotactic sensitivity measured over one segment can be reliably assigned to the background concentration at the midpoint of the segment. Moreover, this time-interval is nearly 1.5 times longer than the average decision making cycle of *Dictyostelium* cells, which has been described as the pseudopod cycle [41]. This means that every CEI measurement is able to represent a gradient sensing and migration process, rather than random changes in cell shape. Because the chemoattractant concentration is known at every point in space, each CEI measurement can be associated with a specific chemoattractant concentration, taken to be the concentration at the midpoint of the corresponding segment.

This gives us an ensemble of independent measurements of track segments, each containing a value for a single cell’s chemotactic efficiency at a known chemoattractant concentration and gradient. Every chemotaxis condition was imaged on at least three different days, and multiple viewing bridges were examined together on each day. For any chemotaxis condition, all the CEI measurements can be binned according to the chemoattractant concentration and a dose response of chemotactic sensing against concentration can be obtained (Fig. 1C).

In total, each chemotaxis condition contributes approximately 50,000 measurements for chemotactic sensing, resulting in more than 2 million measurements throughout the data set (Fig. 2). Together, our pipeline therefore generates an unprecedented ensemble of precisely characterized chemotaxis data that will allow quantitative analysis of chemotactic sensing with real statistical precision.

**FIG. 2.**
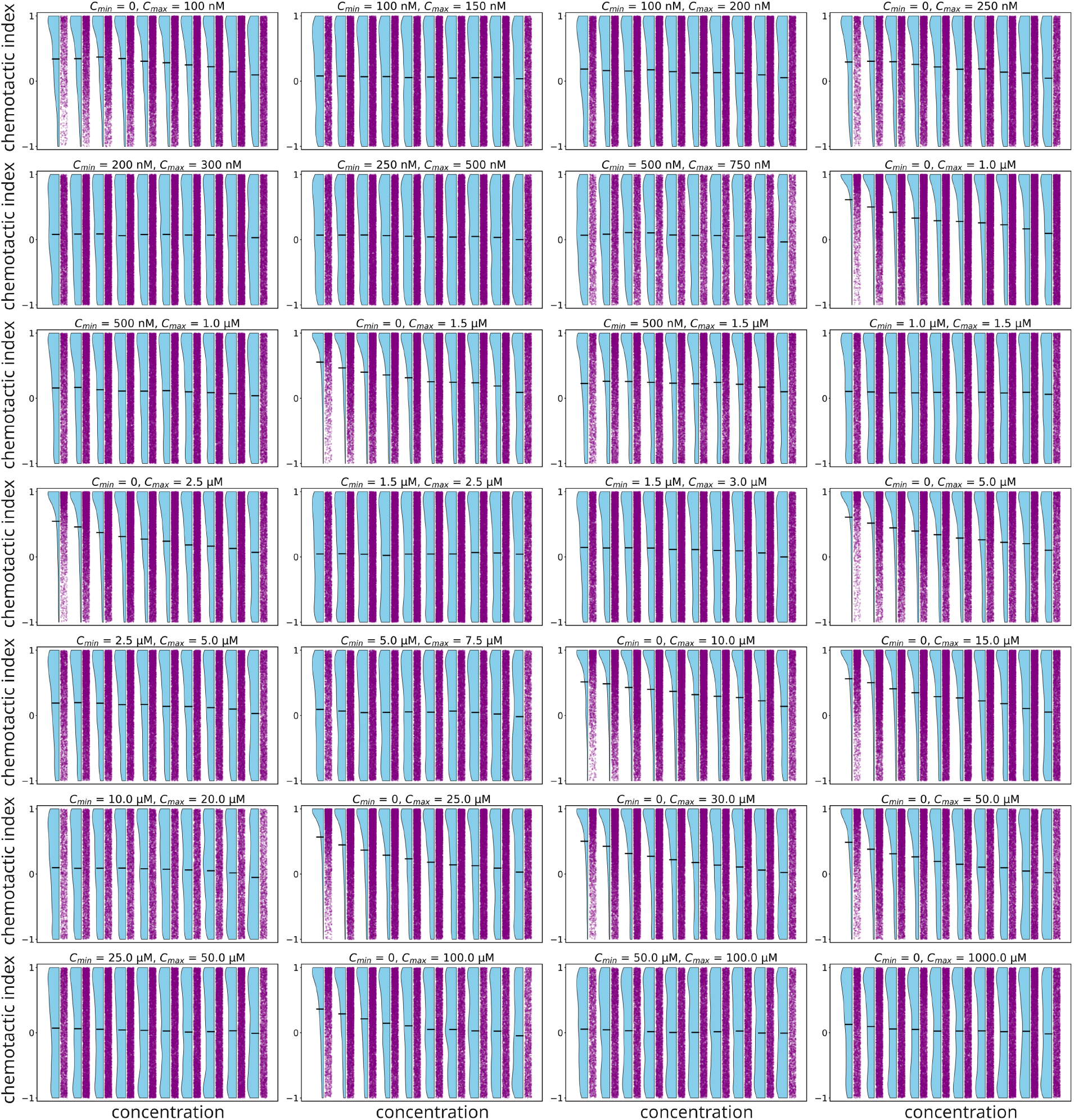
Summary of the ChemoTrack dataset. Chemotactic index (CEI) as a function of concentration for the chemotaxis conditions included in the dataset. Similarly to fig. 1C, each panel shows a gradient with the start and end concentrations in the label, aggregated from 3 independent experiments. Every CEI measurement is associated with a chemoattractant concentration, and all measurements from a chemotaxis condition are are divided into 10 bins uniformly distributed between C_min_ and C_max_, and the mean chemoattractant concentration corresponding to each bin is shown on the x-axis. For each concentration bin, individual CEI measurements are shown as purple dots on the right, the overall distribution is represented by a blue half-violin plot on the left, and the mean CEI is indicated by a horizontal black line. Together with the conditions mentioned in fig. 1C, these comprise all chemotaxis conditions included in ChemoTrack.

### Immediate insights into the mechanism of chemotaxis

We computed the average CEI for each imposed gradient and background concentration, condensing the underlying cell-track data into a quantitative map of chemotactic sensing in the phase space of the imposed gradient and chemoattractant concentration. We assumed instantaneous equilibration of chemoattractant-receptor binding, such that the receptor occupancy, or the fraction of activated receptors in a cell can be assumed to be equal to *C/*(*C* + *K*_*d*_), where *C* is the chemoattractant concentration, and *K*_*d*_ is the dissociation constant for the receptor-ligand binding, which is equal to 1000nM for SpCAMPS binding to cAR1 [33]. Considering the chemoattractant concentration at the front of the cell to be *C*_*f*_, and at the back to be *C*_*b*_, we define the relative difference in chemoattractant concentration across a cell as (*C*_*f*_ − *C*_*b*_)*/C*_*m*_ where *C*_*m*_ is the mean of *C*_*f*_ and *C*_*b*_, representing the average chemoattractant concentration experienced by the cell, also referred to in this paper as the background chemoattractant concentration, and the corresponding background receptor occupancy is equal to *C*_*m*_*/*(*C*_*m*_ + *K*_*d*_). Here the front-rear axis of a cell is taken to be along the direction of the imposed chemoattractant gradient, and the cell-diameter is taken to be 10*µm*, which is approximately equal to the diameter of an unpolarized *D. discoideum* cell.

In fig. 3A we plot CEI as a function of background receptor occupancy and percentage relative difference in chemoattractant concentration between the front and rear of the cell. Considering meaningful chemotactic response to correspond to non-zero values of CEI, we find that our dataset covers the entire region of parameter space associated with a detectable chemotactic response.

**FIG. 3.**
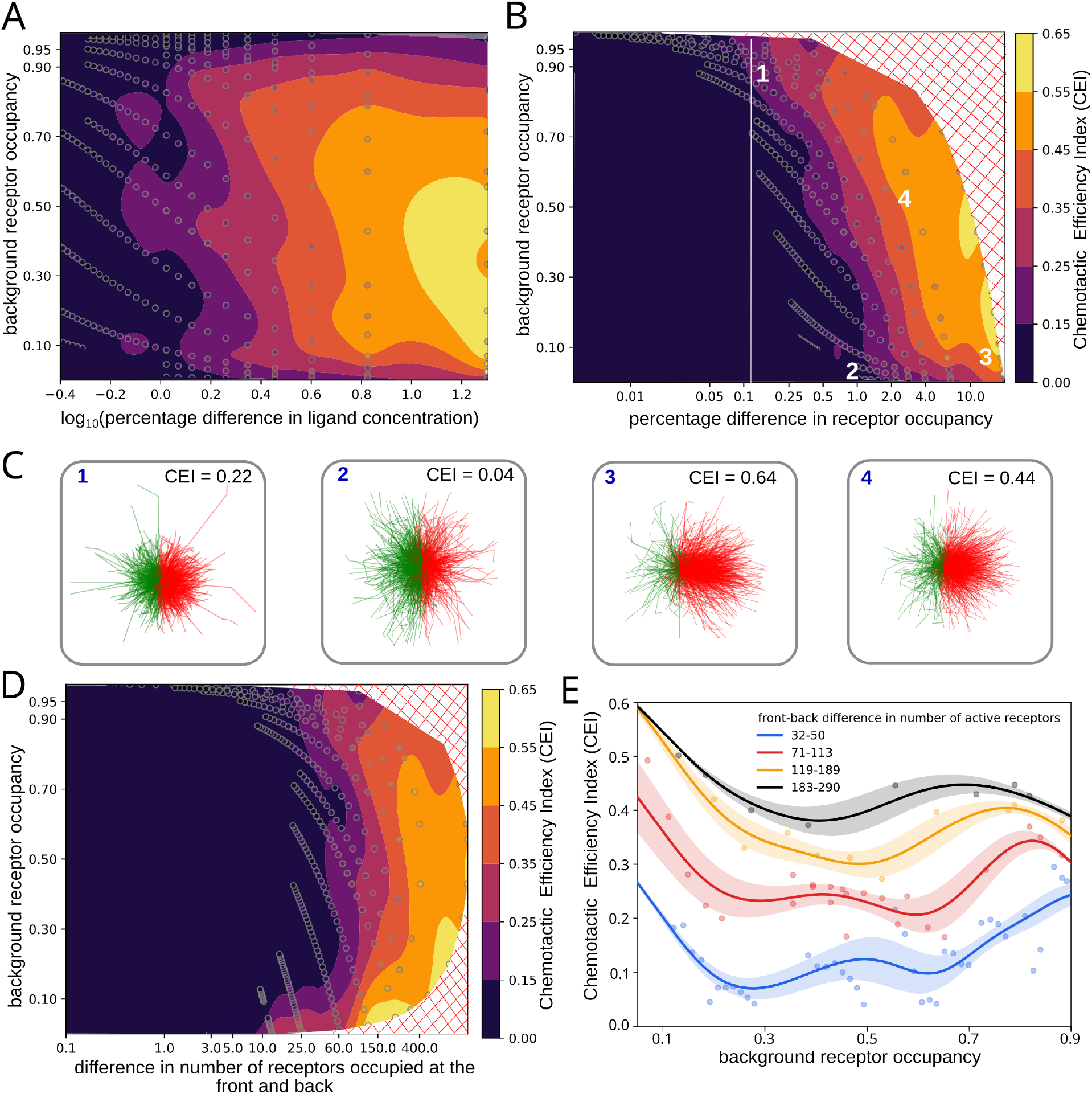
Analysis of chemotactic sensing. (A) Chemotactic efficiency index (CEI) as a function of background receptor occupancy and difference in chemoattractant concentration across a cell. The scatter points denote the data obtained by processing the cell tracks in ChemoTrack, using the method outlined in the previous section. The CEI values at all other points were obtained using radial basis function interpolation with a multiquadric kernel. (B) Re-plotting the data shown in A, as a function of the difference in receptor occupancy across the front and rear of the cell, and the background receptor occupancy. (C) Spider-plots for four points in parameter space that are labeled in B. (D) Re-plotting the data shown in A and B, as a function of the difference in the number of receptors across the front and rear of the cell, and the background receptor occupancy. In panels B and D, the red-hatched region denotes areas of parameter space that are experimentally inaccessible using our experimental design. (E) Chemotactic efficiency index as a function of background receptor occupancy for different values of difference in the number of receptors between front and rear of the cell, obtained by averaging the interpolated CEI over vertical strips in panel D. The dots are the actual data points, the solid line denotes the mean within each range that is obtained from the interpolation, and the shaded region denotes the standard deviation. For generating the dataset, we ignored chemotaxis conditions that result in a chemotactic response that is lower than the threshold for meaningful chemotaxis (regions to the left of the contour boundary for CEI ≈ 0 in A,B, and D). To render the figures, we set these regions to a CEI equal to 0.

Using fig. 3A, we defined the limit of chemotactic sensing to correspond to the contours of CEI = 0, and we can observe that the contour roughly aligns with a one percent difference in chemoattractant concentration, in agreement with a widely-cited limit for chemotactic sensitivity [36].

We defined the fraction of occupied receptors at the front and back as *R*_*f/b*_ = *C*_*f/b*_*/*(*C*_*f/b*_ + *K*_*d*_), where *R*_*f*_ and *R*_*b*_ denote respectively the fraction of active receptors at front and back of the cell. To explore the dependence of chemotactic efficiency on the gradient in receptor occupancy, we plotted our data in fig. 3A by replacing the *x*-axis with the percentage relative difference in receptor occupancy across a cell, and found the limit of chemotactic sensitivity to be much lower than one percent (fig. 3B). In the existing literature, the limits of chemotactic sensing has been widely cited to be equal to a one percent difference in receptor occupancy [42], but our data shows that the limits of sensitivity can be much lower than this. In particular, we observed a meaningful chemotactic response even for a 0.1% difference in receptor occupancy. To visualize this clearly, we compared the cell trajectories at the limit of chemotactic sensing (point 1) with representative examples corresponding to insignificant (point 2), and strong chemotactic responses (points 3,4) (figure 3C).

From fig. 4B, it is clear that cells exhibit markedly different chemotactic sensitivities at different background receptor occupancies despite experiencing the same front–back receptor occupancy difference. This suggests that the receptor occupancy difference alone is insufficient to quantify the chemotactic signal perceived by cells. To explore alternative metrics, we plotted the same data by replacing the *x*-axis with the difference in the number of active receptors between the front and rear of the cell (Fig. 3C), assuming that the number of receptors is equal to 50,000 and equally distributed between the front and rear halves of the cell [43]. On comparing figs. 3B and 3D, we find that the contours for CEI (especially low values of CEI) are aligned better with the *y*-axis in fig. 3D, as compared to fig. 3B, indicating that the limits of chemotactic sensing are more closely governed by the difference in absolute number of active receptors, rather than the difference in the fraction of active receptors. To validate this quantitatively, we performed a regression analysis of the data in figs. 3B and 3D, and found the difference in the number of receptors to be a better predictor of chemotactic sensing (see Supplementary Materials, fig. S4).

**FIG. 4.**
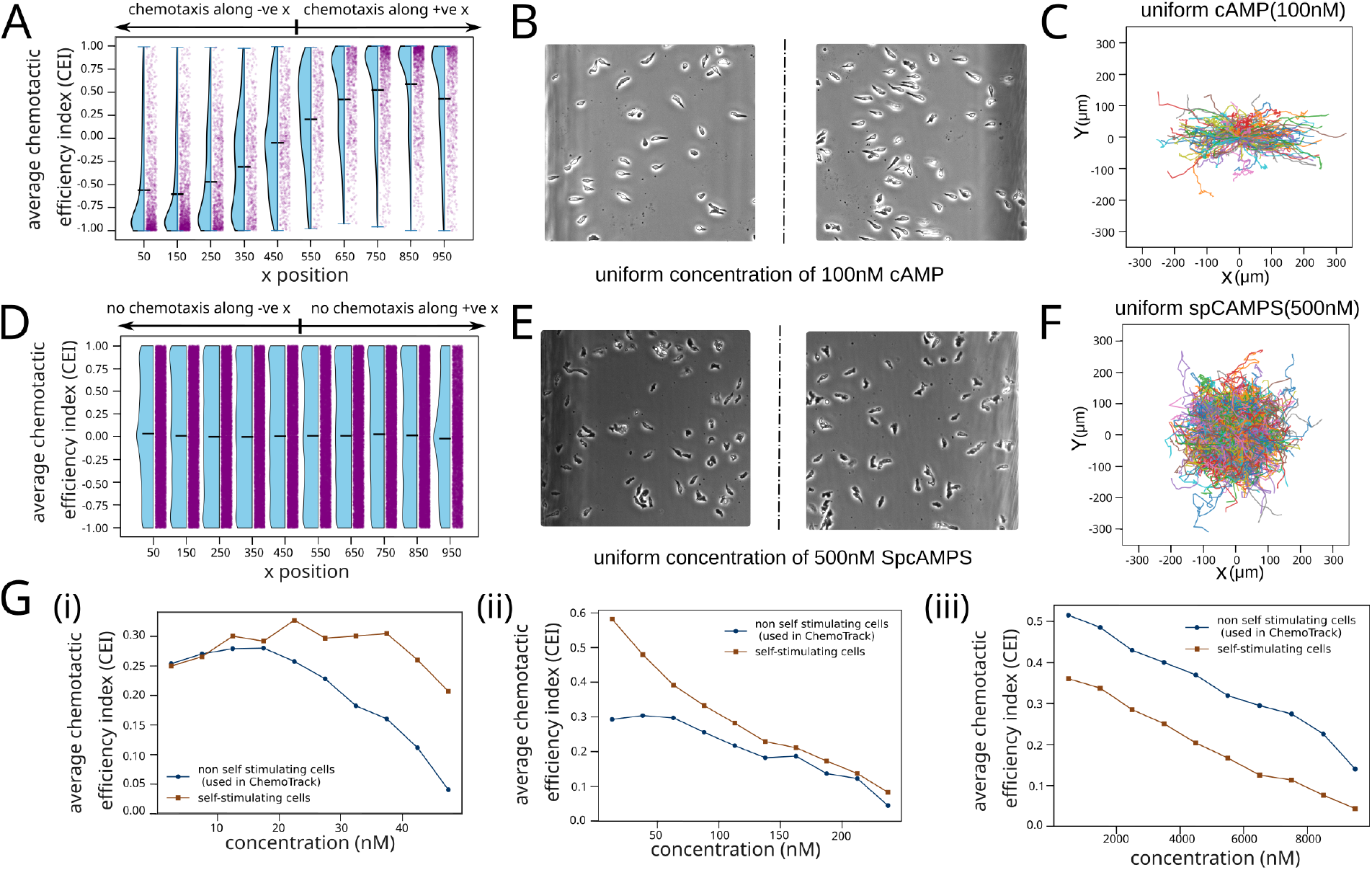
Quantification of the effect of chemoattractant degradation and self-secretion on chemotactic sensing. (A) Chemotactic efficiency index (CEI) as a function of position for a uniform concentration of cAMP equal to 100nM. (B) Snapshots near two ends of the chamber for uniform cAMP concentration, showing cells to be polarized along opposite directions at the two ends. (C) Overlay of individual cell trajectories in uniform 100 nM cAMP, with all tracks aligned to a common starting point (origin), and coloured randomly. (D) CEI vs. position for a uniform concentration of Sp-CAMPS, equal to 500nM. (E) Snapshots near two ends of the chamber for uniform Sp-cAMPS concentration, showing cells to be randomly polarized at all locations. (F) Overlay of individual cell trajectories in uniform 500 nM Sp-CAMPS, with all tracks aligned to a common starting point (origin), and coloured randomly. (G) Comparison of the chemotactic response of WT NC4 and acaA^−^ cells in a gradient of (i) 0-50 nM, (ii) 0-250 nM, and (iii) 0-10 *µ*M Sp-cAMPS.

In the classical chemotaxis literature, the chemoattractant concentration corresponding to maximal chemotactic response has been described to be equal to the *K*_*d*_ of the receptor-ligand binding [36], where half of the receptors are activated, resulting in a background receptor occupancy equal to 0.5. To test this, we plotted CEI with background receptor occupancy, and found that the maximum chemotaxis does not occur at 0.5 (fig. 3E). Instead, we observe that the chemotactic efficiency initially decreases with increase in background occupancy and then increases with a further increase in receptor occupancy. While the initial decrease can be explained by noise in receptor-ligand binding [44], the non-monotonic variation of chemotactic efficiency with background receptor occupancy has not been observed before, and cannot be explained by existing models for eukaryotic chemotaxis. Taken together, the results presented in this section demonstrate that a preliminary analysis of the dataset can be used to validate and test hypotheses connecting statistics of receptor activation with chemotactic sensing.

### Quantifying the effects of degradation and self-secretion

For the data presented in ChemoTrack, cells can neither produce nor degrade the chemoattractant, allowing us to have precise knowledge of the chemotaxis conditions at all points in the domain. Since existing experimental quantification of chemotaxis was performed using either self-stimulating cells or a degradable chemoattractant, we hypothesized that nearly all existing quantitative datasets on chemotactic sensing must be confounded by the effects of chemoattractant secretion and/or self-secretion.

To quantify the effects of degradation, we used the chemoattractant cAMP which can be degraded by the cells, and observed what happens if we have a uniform concentration of cAMP everywhere in the device. We found that even in the absence of an imposed gradient, the cells migrate directionally, and in opposite directions in the two halves of the domain (Movie S1). To quantify this, we plotted the mean CEI as a function of position (Fig. 4A). This clearly shows the cells in the two halves of the domain to be exhibiting a chemotactic response along opposite directions even though there is no imposed chemoattractant gradient. This observation is further supported by representative snapshots of the chamber, in which cells near the two ends are preferentially oriented in opposite directions along the *x*-axis (Fig. 4B). Consistently, an overlay of individual cell trajectories reveals a pronounced bias towards either the positive or negative *x*-direction depending on the cell’s initial position. This is because the cells degrade cAMP everywhere in the chamber, and owing to the geometry of the chamber, the volume of fluid outside the bridge is much greater than the volume inside the bridge. Since the cells are adhered to the cover-slip at a nearly uniform cell density to begin with, the amount of degradation of cAMP per unit volume is much greater in the bridge, leading to an emergent chemoattractant gradient that is directed away from the bridge, resulting in migration of cells out of the bridge. In contrast, cells exposed to a uniform concentration of Sp-cAMPS exhibit a negligible mean chemotactic response (Fig. 4D). Representative snapshots show that cells are randomly oriented throughout the chamber (Fig. 4E), and an overlay of cell trajectories reveals no directional bias, with migration occurring isotropically along all directions (Fig. 4F). These observations validate the effectiveness of the control conditions and demonstrate that chemoattractant degradation can substantially influence the chemotactic response of cells.

To quantify the effects of self-secretion, we used the wild-type (WT) NC4 *Dictyostelium* cells that can secrete cAMP in response to starvation. We quantified the chemotactic efficiency of WT NC4 cells in response to a gradient of Sp-cAMPS and compared it with acaA^−^ cells. For shallow gradients corresponding to a weak chemotactic response, we found that the WT cells exhibit a better chemotactic response (Fig. 4C(i,ii), Movie S2). However, for very steep gradients where the effects of receptor saturation become influential, we found that the WT cells perform worse than acaA^−^ cells (Fig. 4C(iii)). Taken together, the results presented in this section quantify how degradation and self-secretion can significantly alter the response of cells to an imposed chemoattractant gradient.

## DISCUSSION

Like most processes occurring at the intracellular level, the sensing of a chemoattractant gradient by cells is highly stochastic. This is evident in our data, where we found some trajectories to be going nearly opposite to the direction of the imposed gradient even when the majority of them follow the chemoattractant gradient (Fig 1C, middle panel). This data for the first time provides a quantitative measure for the stochasticity in chemotactic sensing by eukaryotic cells. Our data not only quantifies this stochasticity, but also provides accurate estimates for the average chemotactic efficiency of cells, which is lacking in the existing literature. This is because all existing data either used a chemoattractant that can be degraded by the cells, or cells having the ability to secrete chemoattractants, or a combination of both. We showed that a degradable chemoattractant can significantly affect the chemotactic response to an imposed gradient, leading to chemotactic migration in the absence of an externally imposed gradient, or even opposite to an imposed gradient (Movie S3).

Similarly, we showed that the effect of self-secretion of chemoattractants can be complex, either improving the overall chemotactic efficiency or decreasing it, depending on the chemotaxis condition. This means that the effect of chemotactic sensing cannot be inferred by using self-stimulating cells. Since most mammalian cells such as neutrophils [45] and macrophages [46] are known to be self-stimulating, our dataset will serve as a guiding tool for understanding the chemotactic migration of these cells, and isolate the effects of sensing from other relevant features of the chemotactic machinery.

The extensive nature of ChemoTrack enables the testing of different hypothesis for chemotactic sensing. Firstly, we show that a statistical analysis of the data reveals the limits of chemotactic sensing to be more closely related to the absolute difference in number of receptors, as compared to one more widely used metric - the difference in fraction of active receptors [42]. Secondly, using the same analysis, we showed that the chemoattractant concentration corresponding to optimal chemotactic response is not equal to the *K*_*d*_ for receptor-ligand binding. In the existing literature, the underlying mechanism behind *K*_*d*_ being the optimum concentration is that cells are assumed to perform a spatial comparison of the fraction of active receptors, which is proportional to *C/*(*C* + *K*_*d*_), and it can be shown that this comparison is maximally sensitive to change in chemoattractant concentration when *C* = *K*_*d*_ (Fig. S1), implying that a cell will be optimally sensitive to changes in chemoattractant concentration when the concentration is equal to *K*_*d*_. Our data shows that this is not correct, disproving the idea that simple spatial comparison of active receptor numbers as the underlying mechanism for chemotactic sensing in eukaryotic cells.

Furthermore, we found that chemotactic efficiency exhibits adaptation over a wide range of receptor occupancies, with an overall increase at very low and high values of background receptor occupancy. This suggests the presence of supplementary mechanisms for improving chemotactic response in conditions where the chemotactic signal is weak - analogous to the receptor clustering machinery that was discovered by quantitative analysis of bacterial chemotaxis [47]. In the analyses presented in this paper, we have neglected any temporal component to chemotactic sensing, in line with the existing consensus for eukaryotic chemotaxis. However, recent theoretical work suggests that optimal chemotactic sensing for idealized chemotactic agents may involve a combination of spatial and temporal components, depending upon the size, speed, and degree of persistent motility [48] - a prediction that can be tested using our cell tracks. Overall, our dataset - ChemoTrack will be an indispensible tool for quantitative testing and validation of molecular mechanisms and mathematical models for eukaryotic chemotaxis.

## METHODS

### Cell culture and development

We used the *Dictyostelium* NC4 adenylyl cyclase knockout (acaA^−^) cell line detailed in a previous study [31]. *Dictyostelium* cells were grown in lawns of *Klebsiella aerogenes* bacteria on SM agar plates (Formedium). A bacterial suspension was prepared by growing a single bacterial colony in LB media (Formedium) overnight in a shaking-bed incubator at 37^*o*^*C*, and the bacterial lawn was prepared by uniformly spreading 200 *µl* of bacterial suspension on a SM agar plate. cells were scraped from a feeding front and resuspended in 1ml of bacterial suspension. The resulting cell suspension was further diluted in bacterial suspension and approximately 200*µl* of the cell suspension was plated on SM agar plates and cultured in a incubator for 48 hours at 21^*o*^C.

As the *Dictyostelium* cells proliferate, they consume the bacteria ultimately clearing the plate of bacteria. This results in cell starvation that starts their differentiation program, leading to the expression of the chemotaxis receptor cAR1. Since starvation in an agar plate is highly stochastic, it can lead to a large heterogeneity in the chemotactic ability of cells, ultimately leading to inconsistent measurements. We therefore harvest cells from a plate where the bacterial lawn has not started to clear.

The cell suspension was washed in KK_2_ buffer (K phosphate pH 6.2) to remove the bacteria. They were then suspended in 10ml of KK_2_ buffer such that the cell density is 2.5 *\** 10^7^ cells/ml and the cell suspension was continuously shaken to prevent cell anoxia. After 1 hour of starvation, the cells were treated with 100nM cAMP at 6 minute intervals for a duration of 4 hours, while still being shaken. Each chemotaxis condition was performed in triplicate, with each replicate initiating from an independent frozen stock.

### Chemotaxis assay

After 4 hours of development, 1ml of cell suspension was collected, centrifuged and the supernatant containing developmental buffer was removed. The cells were then re-suspended in KK_2_ buffer supplemented with 2 mM MgCl_2_ and 0.2 mM CaCl_2_. The cell suspension was diluted to a final concentration of 3 *\** 10^5^ cells/ml and 300*µl* of this cell suspension was seeded on a standard 22*mm*^2^, number 1 coverslip. The cells were given approximately 15 mins to attach onto the coverslip.

We used the Insall chambers for direct visualization of chemotaxis - a sketch of the chambers is shown in Fig. 1A and further details can be found in [34]. Initially, the entire chamber was filled with the lowest concentration of Sp-cAMPS (BIOLOG Life science institute) that was required for the experiment. Then the coverslip was carefully placed on the chamber such that the tips of the outer well are accessible (see Fig. 1A). The excess fluid was blotted to allow the coverslip to adhere to the chamber through capillary pressure. Finally, the solution from the outer well was aspirated and a chemoattractant solution of the required concentration was injected. The chamber was allowed approximately 5 minutes to rest and then thoroughly sealed with valap to minimize loss of fluid due to evaporation. This leads to the establishment of a one-dimensional gradient of chemoattractant going from the inner well to the outer well.

### Imaging and data processing

The chambers were left to equilibrate for 20 minutes after which the cells on the bridges were imaged at 22^*o*^C for one hour, at 30 second intervals, using a Nikon Ti2 microscope equipped with a 10x objective and a MOMENT-MONO scientific CMOS camera. We obtained our images as 16 bit tiff files, each having a size of 3200×2200 pixels, and processed them using Fiji [40]. For generating the cell tracks, we first subtracted the background using a rolling ball radius of 11 pixels, on a light background with smoothing disabled. This was followed by calibrating the images such that 2.2 pixels equal to 1 *µm*, and converting the images to 8-bit and tracking using an in-house bubble tracking algorithm that outputs the cell centroid over time for every cell, and saves all the tracks in a single csv file. The parameters of the bubble tracking algorithm that were used to generate the cell tracks are shown in Fig. S2A).

## DATA RECORDS

ChemoTrack is available in BioImage archive (accession number: S-BIAD3674) and is licenced under Creative Commons Attribution 4.0 International (CC BY 4.0). The dataset is organized into different directories, each corresponding to a different condition of imposed chemoattractant gradient. Each directory has a name that indicates the experimental conditions. The format for naming the directories is as follows: *C*_min_ − *< C*_min_ *>* − *C*_max_ − *< C*_max_ *>*, where *C*_min_ and *C*_max_ denote respectively the minimum and maximum concentration of Sp-cAMPS in nanomolar (nM), corresponding to that condition. In all the experiments the width of the viewing bridge, along which the gradient was applied was constant (see fig. 1A) and equal to 1mm. Therefore, the chemoattractant concentration can be obtained at any point in the domain through a linear interpolation between *C*_min_ and *C*_max_. For each condition, the dataset includes raw time-lapse microscopy in the form of 8 bit tif files where 1 pixel corresponds to 0.91 *µm*, as we binned our original raw data to enable ease of access, and the original high-resolution images will be made available upon request. In addition, the dataset provides the corresponding cell-tracking data for each replicate in both .csv and .json formats.

## USAGE NOTES

The primary application of this dataset is to support the development of new mathematical models of chemotaxis. In addition to this, the data can be used for training deep learning models for probing whether cell morphology or tracks have any hidden dependence on the chemoattractant gradient, and whether it is be possible to learn the local chemoattractant concentration and gradient experienced by the cell [49]. The tracks are already provided in the dataset, and cell morphology can be extracted from the tiff files using the pre-trained cp-SAM model (Fig. S2B) that is included in the cellpose framework [50], achieving a high level of accuracy even without additional training (Table I).

**TABLE 1.** Results of a cellpose cp-SAM model considering 10 images from different chemotaxis conditions, obtained from our original uncompressed images. For the training, we used five images taken from different chemotaxis comditions and used the default training parameters.

| Model | Precision | Recall | Jaccard | F1-score |
| --- | --- | --- | --- | --- |
| Untrained | 0.9660 | 0.9691 | 0.9373 | 0.9674 |
| Trained | 0.9827 | 0.9977 | 0.9805 | 0.9901 |

## CODE AVAILABILITY

All computer code used to generate the data presented in this paper are freely available in https://github.com/dpp98/Chemotrack--data-analysis-software.

## AUTHOR CONTRIBUTIONS

D.P.P. performed the experiments and analyzed the data. N.S. performed the initial experiments. L.M. and R.H.I. supervised the experimental work. D.V. and P.P. provided inputs on the data analysis. L.T. developed the tracking software. L.M.M. provided useful inputs on the manuscript. R.H.I. conceived the idea for the dataset and supervised the project. All authors revised the paper.

## ACKNOWLEDGMENTS

D.P.P. is grateful to Dr. Abhimanyu Kiran for sharing schematic sketches of the Insall chamber. R.H.I. was supported by grants from the Medical Research Council (MR/X000702/1) and Wellcome Trust (221786/Z/20/Z). P.P. was supported by a UK Research and Innovation (UKRI) Future Leaders Fellowship (MR/V022385/1). L.M.M. was supported by a grant from Cancer Research UK DRCRPG-Nov22/ 100017. The authors are grateful to Prof. Rob R. Kay for his valuable suggestions on the manuscript.

