## Supplementary Text for "ChemoTrack: A comprehensive dataset linking single-cell migration trajectories to precisely defined chemotactic signals"

This PDF file contains Figures S1-S2, Supporting Text, and Legends for Movies S1-S3

### Additional Figures

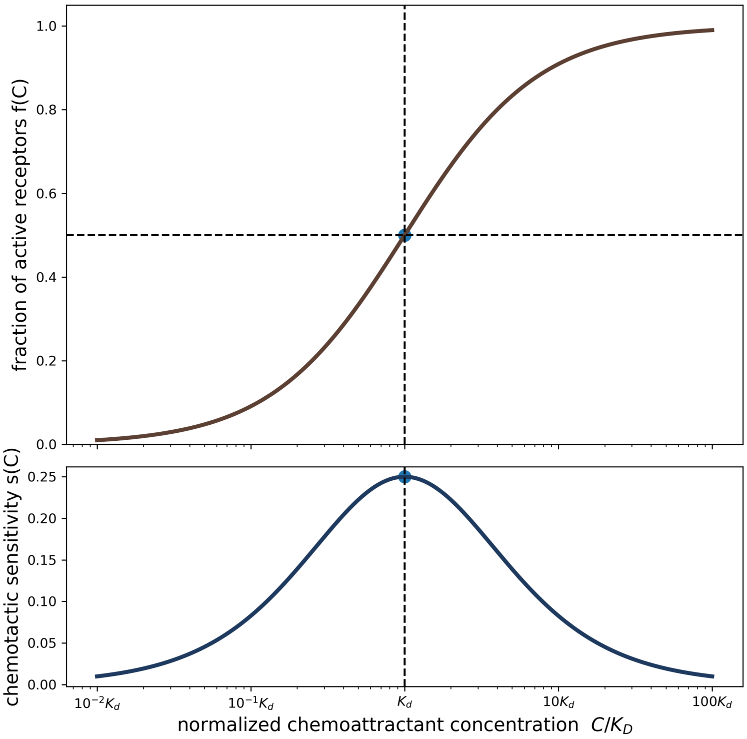


Figure S1: **Chemotactic sensing predicted by a spatial comparison of the active receptors** In the top panel we plot the fraction of active receptors with the chemoattractant concentration, where the fraction of active receptors is given by $f(C) = C/(C + K_d)$, where $C$ and $K_d$ are respectively the chemoattractant concentration and the dissociation constant for the receptor-ligand binding. Assuming that cells respond to chemoattractant gradients by detecting fractional changes in the receptor occupancy, the sensitivity is given by $s(C) = \frac{df}{\frac{dC}{C}} = C\frac{df}{dC} = \frac{CK_d}{(C+K_d)^2}$. We plot this quantity in the bottom panel and show that the maximum sensitivity is obtained when the chemoattractant concentration is equal to $K_d$.

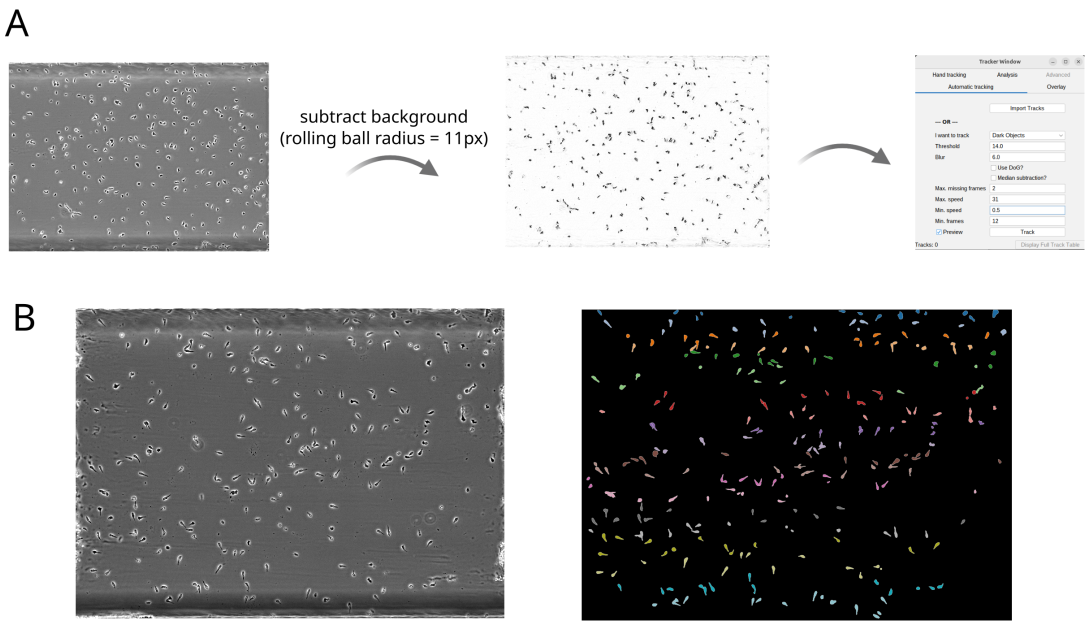


Figure S2: (A) Schematic of the segmentation and tracking of cells using our inhouse FiJi plugin. (B) Segmentation of the cell boundaries using the untrained cp-SAM model in cellpose-v4.

### Movies

#### Movie S1. Cells in uniform cAMP

Time-lapse video showing acaA$^{-}$ cells exposed to a uniform 100nM of the degradable chemoattractant cAMP. We can observe that even in the absence of an externally imposed chemoattractant gradient, cells can migrate directionally due to self-generated chemoattractant gradients, and this directional response is quantified in Fig. 3A of the main text.

#### Movie S2. Effect of self-secretion of chemoattractants on chemotactic migration

Comparison of two time-lapse videos showing identical, shallow chemoattractant gradients of 0-250nM Sp-cAMPS (higher to the right). Left panel shows the acaA$^{-}$ cells that are used for generating ChemoTrack. Right panel shows corresponding wild-type NC4 cells that are identical, except that they express the acaA gene and are therefore able to secrete cAMP in response to starvation.

#### Movie S3. Chemotaxis in presence of a degradable chemoattractant

Time-lapse video showing cells responding to a gradient of 50nM-150nM of cAMP (higher to the right).

### Supporting text

#### S1 Comparison of CEI with other well-known metrics for chemotactic sensing

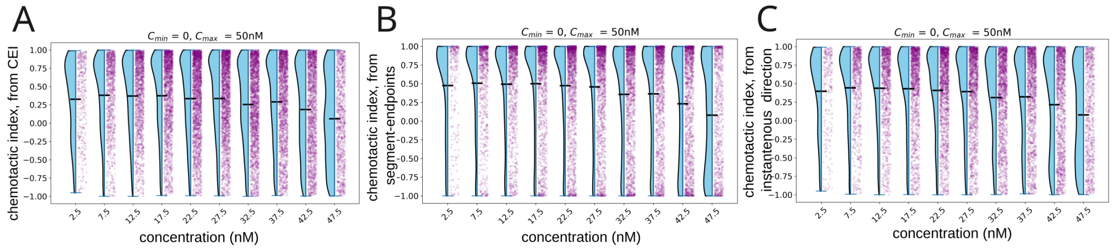


Figure S3: **Comparison of chemotactic sensing from different metrics.** (A) Dose response of chemotaxis with respect to background concentration, computed using the chemotactic efficiency index. (B) Same data with the chemotactic index quantified using the cosine of the angle made by the displacement vector from the initial and final points of a segment. (C) Same data with the chemotactic index quantified by computing the cosine of the displacement vector between two consecutive points in a segment.

To justify the use of CEI as a reliable metric for chemotactic sensing, we compared it with two other metrics to show how the choice of an inappropriate metric for chemotaxis can lead to misinterpretation of data. Some of the commonly used metrics in both in experimental and theoretical quantification of chemotactic sensing rely on computing the cosine of the angle formed by the imposed gradient and the direction of cell motility ($< cos(\theta) >$) [1, 2]. We compared the chemotactic sensing obtained using CEI (fig. S3A), with two other metrics that are based on $< cos(\theta) >$. In fig. S3B, we quantified chemotactic sensing using the cosine of the angle formed between the displacement vector going from the initial to the final point of a segment, and the positive x-axis, and in fig. S3C, we used the average of the cosine of the instantaneous cell orientation obtained from two consecutive points in the cell track. In both the $< cos(\theta) >$ based metrics, we observed a bimodal distribution in the chemotactic index, indicating the presence of a subpopulation of cells exhibiting chemo-repulsive behaviour, that can be seen by the peaks near $y = -1$, which corresponds to migration opposite to the direction of the imposed gradient. However, this is merely an artefact of using an inappropriate metric, and is absent when chemotactic efficiency is quantified using CEI (Fig. S3A).

#### S2 Comparison of difference in receptor number vs. receptor occupancy

In figs. 3B and 3D of the main text, we show CEI as a function of background receptor occupancy and either the front–back difference in the fraction of active receptors (Fig. 3B), or the front–back difference in the number of active receptors (Fig. 3D). To quantitatively test which amongst these two is a better predictor of chemotactic sensing, we performed a multiple linear regression analysis and found that the absolute front–back receptor difference was a strong stand-alone predictor of chemotactic efficiency, explaining approximately 61% of the observed variance in CEI (Fig S3A), with a coefficient of determination $R^2 = 0.615$. Incorporating background receptor occupancy into the model increased the explained variance only marginally (($R^2 = 0.621$)). This indicates that background receptor occupancy provides little additional predictive information once the absolute difference in active receptors is known. In contrast, the relative front–back occupancy difference alone explained only 48% of the variance in CEI (Fig S3B), with a coefficient of determination $R^2 = 0.483$, and including background receptor occupancy substantially improved the model (($R^2 = 0.707$)). This suggests that the front–back difference in the absolute number of active receptors is a better predictor of chemotactic efficiency as compared to the difference in fraction of active receptors.

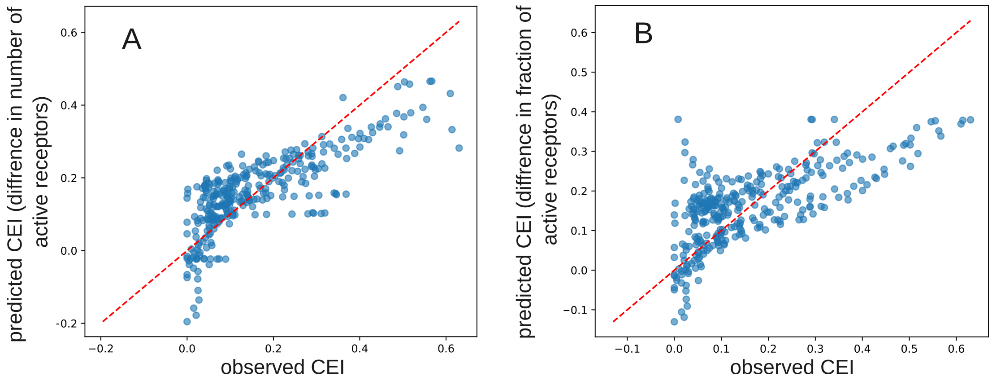


Figure S4: **Regression analysis for receptor activation** (A) Observed versus predicted chemotactic efficiency index (CEI) obtained from a linear regression model using the front–back difference in the number of active receptors as the sole predictor. (B) Observed versus predicted CEI obtained from an analogous model using the front–back difference in the fraction of active receptors as the sole predictor. Each point represents the average CEI obtained from a particular gradient and background chemoattractant concentration, and the dashed red line indicates perfect agreement between predicted and observed values (($y = x$)). Tighter clustering of points around the identity line in A indicates a higher predictive power of the difference in absolute number of active receptors compared with difference in fraction of active receptors.
